# Connectivity of marine predators over the Patagonian Shelf during the highly pathogenic avian influenza (HPAI) outbreak

**DOI:** 10.1101/2023.12.12.570574

**Authors:** Javed Riaz, Rachael A. Orben, Amandine Gamble, Megan Tierney, Paulo Catry, José P. Granadeiro, Letizia Campioni, Alastair M. M. Baylis

## Abstract

Animal movement and population connectivity are key areas of uncertainty in efforts to understand and predict the spread of infectious disease. The emergence of highly pathogenic avian influenza (HPAI) in South America poses a significant threat to globally significant populations of colonial breeding marine predators in the South Atlantic. Yet, there is a poor understanding of which species or migratory pathways may facilitate disease spread. Compiling one of the largest available animal tracking datasets in the South Atlantic, we examine connectivity and inter-population mixing for colonial breeding marine predators tagged at the Falkland Islands. We reveal extensive connectivity for three regionally dominant and gregarious species over the Patagonian Shelf. Black browed albatrosses (BBA), South American fur seals (SAFS) and Magellanic penguins (MAG) used coastal waters along the Atlantic coast of South America (Argentina and Uruguay). These behaviours were recorded at or in close proximity to breeding colonies and haul-out areas with dense aggregations of marine predators. Transit times to and from the Falkland Islands to the continental coast ranged from 0.2 – 70 days, with 84% of animals making this transit within 4 days - a conservative estimate for HPAI infectious period. Our findings show the incursion of HPAI to the Falkland Islands marine predator community is a highly credible threat, which may be facilitated by BBA, SAFS and MAG connectivity with South America. This information is vital in supporting HPAI disease surveillance, risk assessment and marine management efforts across the region.

**Significance:** The recent emergence of highly pathogenic avian influenza (HPAI) in South America poses a major threat to globally significant marine predator populations in the South Atlantic. There is extensive connectivity over the southern Patagonian Shelf between regionally dominant seal and seabird populations, with potential for large-scale pathogen spread. Despite this connectivity, outbreaks of HPAI are unevenly distributed across the region. Connectivity information is integral for regional disease surveillance, predictive modelling and population viability assessments.

## 1 Introduction

Animal movement plays a fundamental role in shaping the ecological and evolutionary dynamics of wild populations (Morales et al. 2010, Sutherland et al. 2013). Spatial connectivity between populations is essential for maintaining genetic diversity and long-term population persistence and viability (Bowler & Benton 2005, Bicknell et al. 2012). However, animal movements can also facilitate the dispersal of pathogens they host (via direct or indirect contact between infected and susceptible individuals), posing a significant threat to population health and conservation status (Altizer et al. 2011). Movement behaviours (i.e. foraging, prospecting, resting and transit) play different roles in disease transmission (Boulinier et al. 2016), and characterising these mechanisms is critical to understand how pathogens spread through spatial networks (Daversa et al. 2017).

Since 2021, a highly pathogenic avian influenza (HPAI) panzootic has had dramatic impacts on wild populations around the world, causing mass mortalities in numerous seabird and marine mammal populations (Falchieri et al. 2022, Klaassen & Wille 2023, Leguia et al. 2023, Lane et al. 2023). Driven by connectivity over vast spatial scales, HPAI is spreading at unprecedented rates (Boulinier 2023, Jeglinski et al. 2023, Klaassen & Wille 2023) and is predicted to impact geographically remote, high-latitude marine predator communities in the southern hemisphere, which have traditionally been insulated from periodic HPAI outbreaks affecting wild populations in Eurasia, Africa and North America (Dewar et al. 2022).

The Patagonian Shelf Large Marine Ecosystem (LME) spans the Atlantic coast of South America, encompassing the continental shelf areas adjacent to Uruguay, Argentina and the Falkland Islands. It is regarded as one of the most productive marine ecosystems in the world, supporting a diverse range of colonial breeding marine predators (i.e. pinnipeds, penguins and flying seabirds) (Croxall & Wood 2002, van der Grient et al. 2023). For these animals, the Falkland Islands are one of the most important breeding locations in the Patagonian Shelf LME (Baylis et al. 2019b, 2021). For example, the Falkland Islands are home to approximately 75% of the world’s black-browed albatross (*Thalassarche melanophris*), 50% of the global South American fur seal (*Arctocephalus australis*) population, and globally significant populations of southern rockhopper (*Eudyptes chrysocome*), gentoo (*Pygoscelis papua*) and Magellanic (*Spheniscus magellanicus*) penguins (Baylis et al. 2013b a, 2019a, Wakefield et al. 2014, Falabella et al. 2019).

Despite the relative geographic isolation of the Falkland Islands, HPAI is an emerging threat to its globally significant seabird and pinniped populations (Bennison et al. 2023). Biological risk assessments seeking to identify spatial and temporal components of HPAI spread in the region have been hampered by a limited understanding of animal movement, specifically, population connectivity and migration pathways (Dewar et al. 2023). Uncertainty about which candidate species or spatial networks act as vectors of disease transmission can undermine predictive modelling efforts (Webster et al. 2017), ultimately affecting disease surveillance and the development of adaptive conservation and management strategies (Bestley et al. 2020, Murphy et al. 2021).

In this study, we compile an extensive multi-year telemetry dataset for regionally representative colonial marine predators over the Patagonian Shelf to assess population connectivity and networks for disease transmission. Using data from nine seabird and pinniped species tagged at the Falkland Islands with Platform Terminal Transmitters (PTT) and Global Positioning System (GPS) devices, we quantify residency in adjacent South American coastal waters (Argentina and Uruguay) and identify hotspots of spatiotemporal connectivity between populations. Our findings can support HPAI risk assessments in the region and improve capacity to predict future threats posed by new infectious diseases, guiding surveillance efforts and population viability assessments.

## 2 Methods

In this study, we compiled GPS and PTT satellite telemetry datasets available for nine colonial breeding marine predators at the Falkland Islands. This included 3 pinniped species (South American fur seal; South American sea lion [*Otaria flavescens*] and Southern elephant seal [*Mirounga leonine*]), 4 penguin species (Magellanic; gentoo; southern rockhopper and King [*Aptenodytes patagonicus*]) and 2 species of flying seabirds (Black-browed albatross and sooty shearwater [*Ardenna grisea*]). While light-based geolocation (GLS) tracking data are also available for some penguin and flying seabird species, these data were excluded due to low spatial precision over local and regional scales (see Table S1 for full details of compiled tracking data). Following visual inspection of tracking data from each of the nine species, only Black browed albatrosses (BBA), South American fur seals (SAFS) and Magellanic penguins (MAG) showed evidence of regional-ranging movements to coastal waters in South America. Therefore, we restricted the collated tracking dataset to these three species. Details of the tracking data compiled are provided in Table 1 and Table S1. Animal handling and tag deployment procedures are available in source literature (Campioni et al. 2017, Baylis et al. 2019b, Riaz et al. 2023) and supplementary material.

**Table 1.**
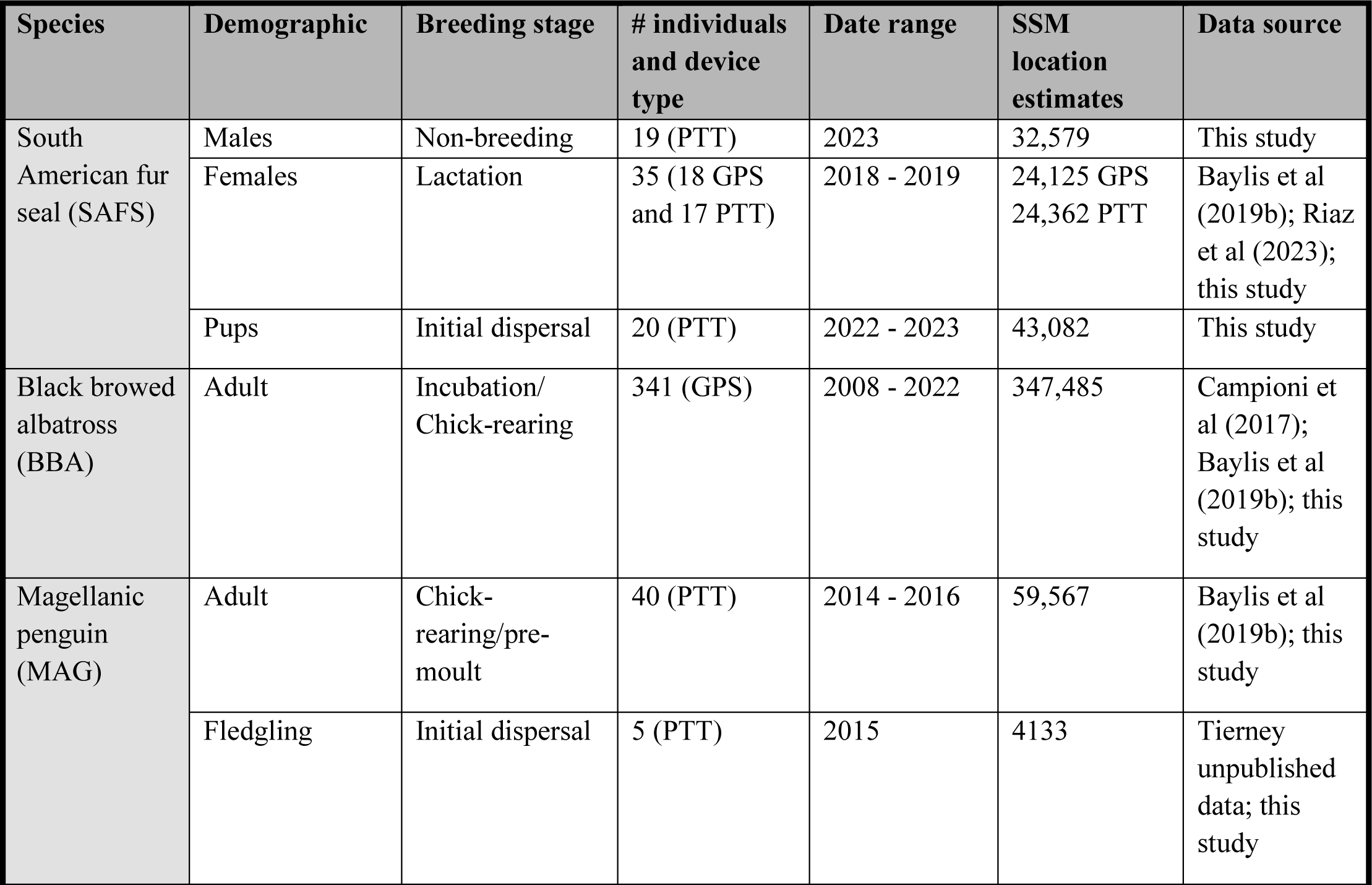
Summary of satellite telemetry data and tracking efforts for the three colonial breeding marine predators at the Falkland Islands. Some BBA and MAG individuals were tracked over multiple life history stages (i.e. incubation/chick-rearing and chick-rearing/moult). The total number of location estimates after state-space model (SSM) data processing are also provided (See *Methods* for details). See Table S1 for details of the number and proportion of colonies tracked.

All data processing and analyses were performed using R statistical software (version 4.3.1; R Core Team 2022). Raw location data for the three species were assembled, plotted and visually inspected. Data were then subjected to quality-control checks adapted from the framework provided in Ropert-Coudert (2020). Near-duplicate location estimates were removed (locations occurring within 2 minutes for PTT data and 10 seconds for GPS data). We also removed locations implying unrealistic travel speeds within movement trajectories (exceeding 4 m s^−^1 for SAFS, 10 m s^−^1 for MAG and 20 m s^−^1 for BBA). With the quality-controlled data for the three species, we fitted a random walk State-Space Model (SSM) using the ‘*aniMotum’* package (Jonsen et al. 2023). This approach provides location estimates at regularised time-steps along movement trajectories, while also accounting for observation errors in tracking data (Jonsen et al. 2023). All PTT data for SAFS and MAG were regularised at 1-hour time steps, while GPS data for SAFS and BBA were regularised at 15-minute and 10-minute time steps, respectively. The programming of these species- and device-specific time-steps within the movement model were chosen based on visual and comparative assessment of predicted location estimates with the raw location fixes for each individual (Riaz et al. 2021, 2023). The final SSM dataset contained location information for 460 individuals, which comprised of 341 BBA, 74 SAFS and 45 MAG. Tracking data was available throughout most of the annual cycle for SAFS, while BBA and MAG tracking data were restricted to the summer breeding and pre-moult period (Fig. S1).

To quantify movement along predicted location estimates and assess where individuals spend disproportionally more or less time, we also fitted a time-varying move persistence (*γ*_*t*_) model to SSM location data (‘*aniMotum*’ package; Jonsen et al. 2023). This indexed changes in movement behaviour as a continuous variable (ranging from 0 – 1) based on autocorrelation in both speed and direction (Jonsen et al. 2023). Relatively low *γ*_*t*_values are indicative of residency (low speed and directionality) in movement trajectories typically encompassing rest or more stationary foraging behaviours. Conversely, relatively high values represent more directed movements (high travel speeds and linear directionality) and encompass transit and larger-scale searching movements (Jonsen et al. 2019, Riaz et al. 2021, Grecian et al. 2022). For each individual, *γ*_*t*_ values were normalised, rescaling all estimates to span the full range (0 through to 1). This approach offers a more detailed and nuanced quantification of changes in behaviour along movement trajectories, and enables comparison between species which move over different spatiotemporal scales (Jonsen et al. 2023).

With the processed movement dataset for BBA, SAFS and MAG, we quantified the degree of connectivity between the Falkland Islands and adjacent South American coastal habitats. For each individual, we calculated the proportion of time spent in connection with the South American coastline. This was defined as any animal location recorded between land and the limit of the territorial sea maritime boundary (12 nautical miles from baseline [∼ 25 km]). This spatial delineation was chosen primarily because it was considered to be a reasonable measure of distance to land where high aggregations of colonial breeding marine predators are likely to occur. As an additional justification, the boundary is also relevant to national marine conservation and management efforts (Kraska et al. 2015). To understand the temporal components of connectivity, we also calculated the time taken for individual animals to travel between the coastal habitats of the Falkland Islands and the South America.

To identify areas where and when individuals were likely to be resident in coastal South America, we used the continuous behavioural index of persistence (*γ*_*t*_) in movement trajectories. The *γ*_*t*_parameter is traditionally used to quantify behavioural states at-sea, with low *γ*_*t*_assumed to represent area-restricted search (ARS) foraging behaviours (Jonsen et al. 2019, Grecian et al. 2022, Shuert et al. 2022). However, areas of low *γ*_*t*_may also represent resting or haul-out (i.e. land-based resting for pinnipeds) behaviours (Riaz et al. 2021, Thums et al. 2022, Oosthuizen et al. 2022). We defined land-associated residency behaviours as any location either over land or within 1 km of land where normalised *γ*_*t*_values were < 0.5. Our approach summarising *γ*_*t*_ into a binary measure was considered pragmatic to calculate discrete land-associated behavioural states (Thums et al. 2022). Finally, the individual tracks reaching the coast were inspected for evidence of haul-out or roosting behaviour (e.g. Kralj et al. 2023) To assess evidence of intra-specific and inter-colony interactions across the Patagonian Shelf LME, spatial information of all known BBA, SAFS and MAG breeding and haul-out sites in the region were collated from published sources (Table S2) (Crespo et al. 2015, Phillips et al. 2016, Franco-Trecu 2017, Dodino et al. 2022, Garcia-Borboroglu et al. 2022, Millones et al. 2022, Raya Rey et al. 2022).

## 3 Results

For each species, multiple individuals were distributed in close connection with South American breeding/haul-out sites where conspecifics and other colonial marine predators congregate (Fig. 1). While transit times to and from the Falklands Islands - South American coast took up to 70 days for some individuals, the vast majority (84%) made this trip within 4 days. Notably, individuals of BBA, SAFS, and MAG that migrated to coastal South America also made return trips to the Falkland Islands. While the number of individuals transiting between coastal waters is influenced by the coverage of the tracking data, we found connectivity occurred throughout the annual cycle (Fig. S1).

**Fig. 1.**
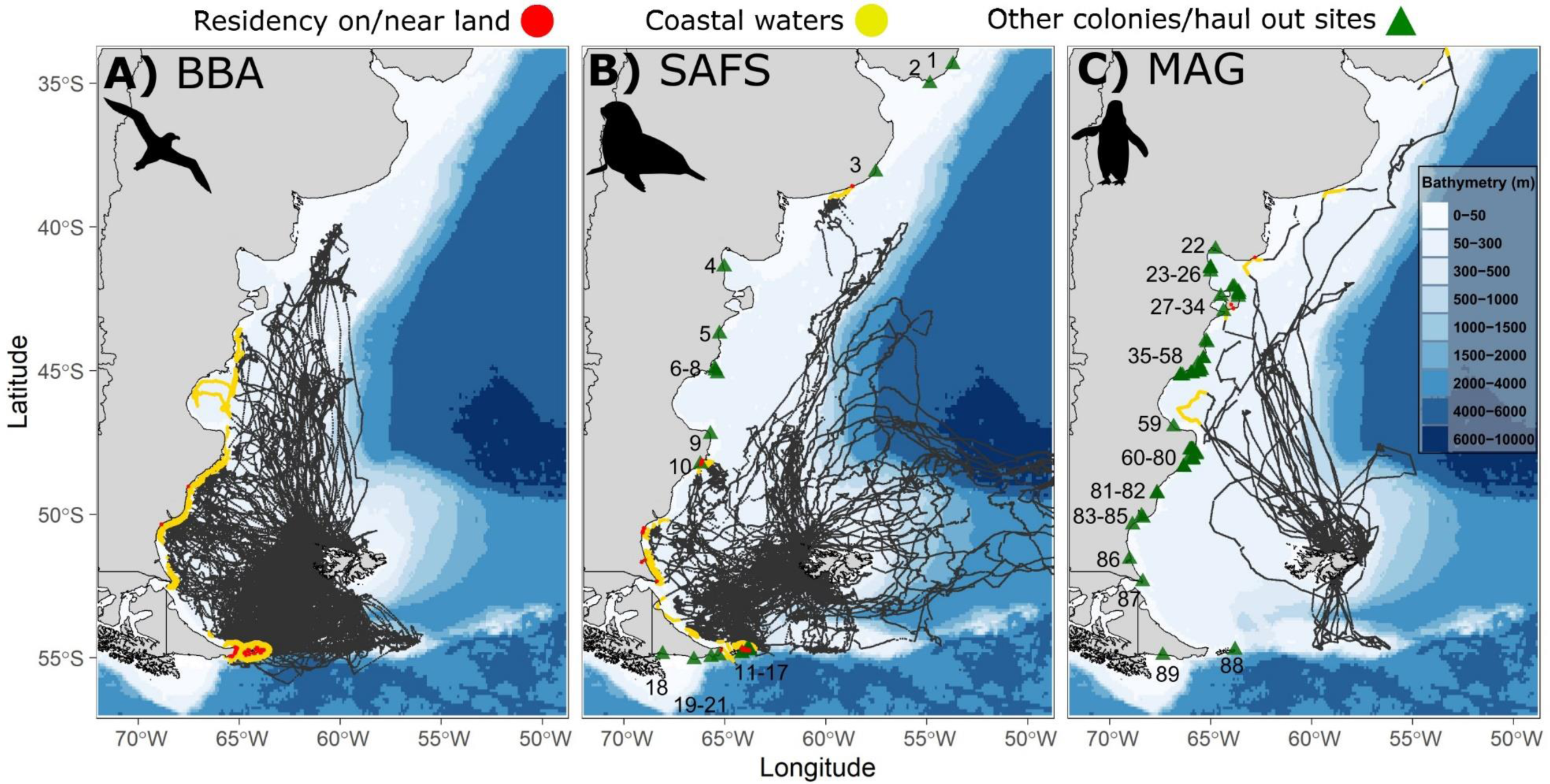
State-space model (SSM) location estimates for BBA (n = 341), SAFS (n = 74) and MAG (n = 45) individuals tracked at the Falkland Islands (see *Methods* for details). Dark grey points represent all SSM locations estimates outside the coastal waters of Argentina and Uruguay, delineated as the boundary between land and the limit of the territorial sea (12 nautical miles from baseline). Yellow points indicate all location estimates within this coastal boundary, while red points illustrate residency on/near land (see *Methods* for details). Green triangles (and associated numbers) summarise other known breeding or haul-out sites across the southern Patagonian Shelf region for these three species (see Table S2). Bathymetric gradients displayed were created using the ‘*ggOceanMaps’* package (Vihtakari 2022).

Black browed albatrosses displayed a high degree of connectivity (32% of individuals) extending along the east coast of Argentina from Staten Island to Peninsula Valdes (approximately -55° to -44°S) (Fig. 1). These individuals spent an average of 13.6% of their trips (36.7 hours) in Argentine coastal waters (means and 95% CI provided in Table 2). Connectivity was particularly pronounced in waters surrounding Staten Island, which hosted 88 BBA individuals (81% of all individuals exhibiting connectivity). Most individuals performed several foraging trips between the Falkland Islands and South America, with transit times lasting < 5 days in duration (Fig. 2). Staten Island was also an important area of residency for 25 (28%) of these individuals. While we did not find any evidence of BBA roosting on land in South America, individuals displayed nearshore rafting or low move persistence over land. Two BBA individuals also recorded these behaviours slightly further north along the Argentine coast (-52° to -49°S) (Fig. 1). The mean duration individuals spent in nearshore residency was 0.8 hours (0.4% of trips), although this lasted up to 5 hours for some individuals in Argentina (Table 2).

**Fig. 2.**
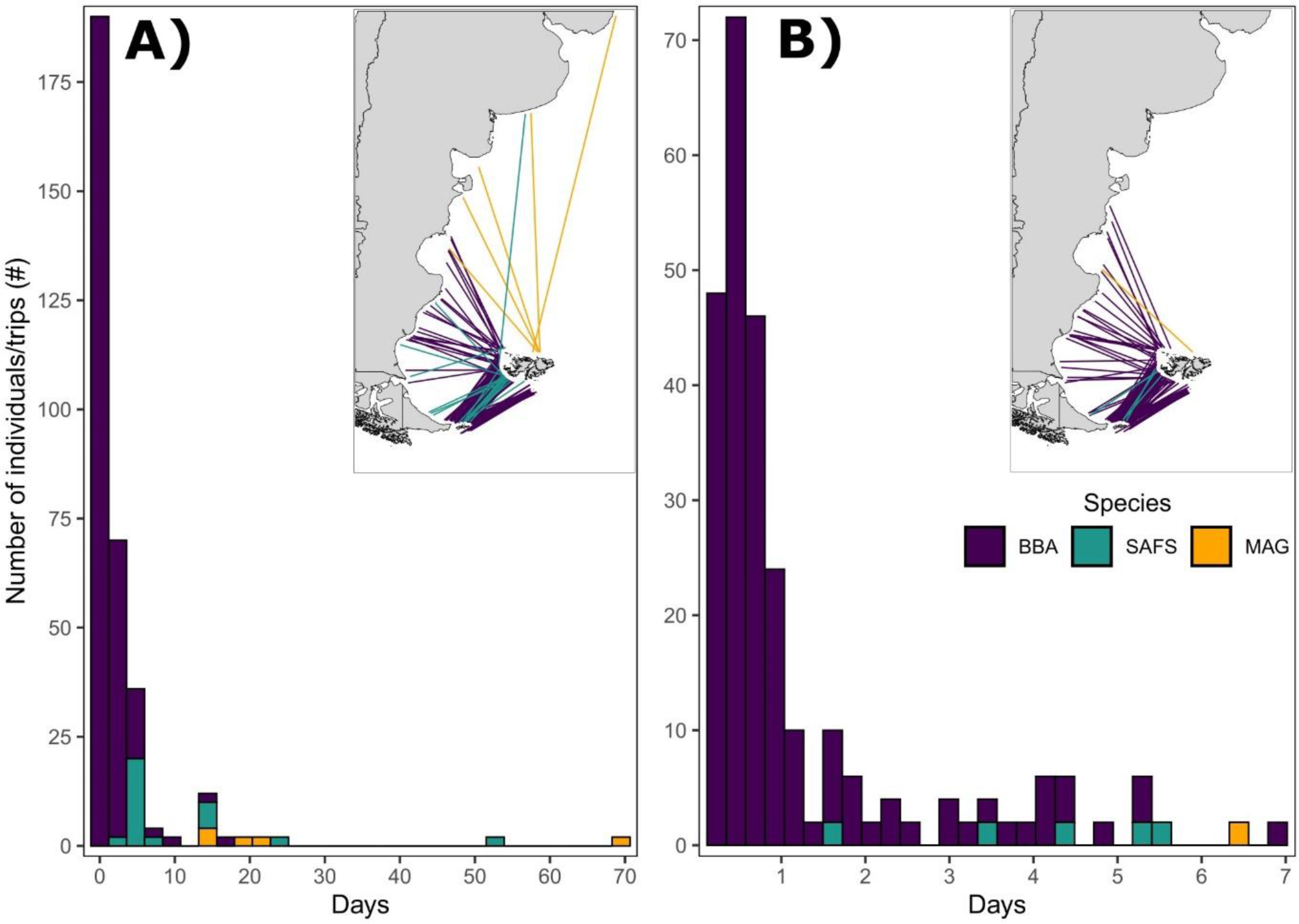
Time taken for individuals to travel between A) the Falkland Islands and South America during outbound migrations, and B) from South America to the Falkland Islands during inbound migrations. These travel times are calculated based on the last and the first location recorded in coastal waters (territorial sea) during inbound and outbound trips. Corresponding colours for BBA, SAFS and MAG are presented in the histogram and embedded map.

**Table 2.** Summary of the proportion (%) and amount (hours) of time each species (BBA, SAFS and MAG) spent in South American coastal waters and in resident behaviours on/near land. Mean (95% confidence intervals [CI]) and range values are provided, calculated across individuals which recorded connectivity with South America.

|  | Coastal waters |  | Residency on/near land |  |
| --- | --- | --- | --- | --- |
|  | Mean (95% CI) | Range | Mean (95% CI) | Range |
|  | **BBA (n = 109)** |  | **BBA (n = 27)** |  |
| Proportion of time (%) | 13.6 (10.9 – 16.3) | 0.1 – 67.8 | 0.4 (0.1 – 0.8) | 0.03 – 5.4 |
| Total time (hours) | 36.7 (27.3 – 46.0) | 0.2 – 298.9 | 0.8 (0.38 – 1.2) | 0.17 – 5.0 |
|  | **SAFS (n = 17)** |  | **SAFS (n = 13)** |  |
| Proportion of time (%) | 21.6 (12.9 – 30.2) | 3.7 – 67.8 | 4.2 (1.1 – 7.3) | 0.1 – 17.6 |
| Total time (hours) | 365.9 (181.7 – 550.2) | 26.0 – 1268.3 | 76.9 (19.6 – 134.2) | 1.0 – 329.8 |
|  | **MAG (n = 5)** |  | **MAG (n = 2)** |  |
| Proportion of time (%) | 12.9 (0.2 – 25.7) | 1.6 – 28.1 | 2.9 (0.1 – 37.5) | 0.2 – 5.6 |
| Total time (hours) | 104.1 (0.1 – 214.0) | 29.0 – 241.8 | 14.5 (0.1 – 185.7) | 1.0 – 27.9 |

Of all SAFS individuals tracked, 17 (23%) were recorded in coastal Argentine waters (Fig. 1; Fig 2). These 17 individuals comprised of all demographic groups examined (males, females and pups) (Fig. S2). The mean proportion of time spent in Argentine coastal waters was 22% (366 hours) (Table 2). Locations in Argentine coastal waters were primarily concentrated (n = 14 individuals) around the breeding/haul-out sites at Staten Island (Fig. 1 icon 11 – 17; Table S2), but also extended northwards along the coastline to the SAFS haul-out at Islote del Cabo (Fig. 1 icon 10; Table S2). Inbound/return migrations from South America to the Falkland Islands were recorded for both breeding females and non-breeding males, and generally took between 1 -5 days (Fig. 2). Most of the SAFS recorded in this region of southern Argentina also hauled-out on land (n = 12), with 10 males hauling out on Staten Island (in multiple locations), 1 male hauling out at Islote del Cabo and 1 pup hauling out at multiple locations between these sites (-54° to -51°S) (Fig. 1; Fig. S2). One male was also recorded resting in northern Argentinian waters, near the haul-out site at Mar Del Plata (Fig. 1 icon 3; Table S2). On average, haul-out periods lasted 77 hours in duration, although one male was hauled-out for over 13 days on Staten Island (Table 2).

For MAG, 5 individuals (3 fledglings and 2 adults; 11% of all individuals) were recorded in close proximity to South America (Fig. 1). On average, these individuals spent 13% (104 hours) of their trip in these coastal areas (Table 2). All 5 MAG individuals were recorded nearby to other MAG breeding sites (Fig. 1 icons 22 – 80; Table S2) in coastal regions of Argentina, and 2 individuals ventured into more northern waters located in Argentina and Uruguay. Two individuals were recorded resting on land in northern Argentina, with one at Peninsula Valdes for over 1 day in duration (Fig 1 icons 27-34; Table S2). One MAG adult returned from coastal Argentine waters to the Falklands Islands in ∼ 6 days (Fig. 2).

## 4 Discussion

This study is the first to quantitatively assess spatiotemporal connectivity of colonial marine predators over the Patagonian Shelf. By compiling the largest dataset available for marine predators tagged at the Falkland Islands to date, we reveal a high degree of regional connectivity and strong evidence for inter-population mixing for BBA, SAFS and MAG – three highly abundant and gregarious colonial marine predators within the Patagonian Shelf LME. Our findings significantly improve understanding of the candidate species and spatial networks which may facilitate the spread of HPAI and other infectious diseases from South America. This information is critical in developing evidence-based risk assessments capable of evaluating geographic vulnerabilities and predicting disease spread across space and time.

In this study, we show Falkland Island populations of BBA, SAFS and MAG have extensive connectivity with adjacent South American coastal habitats. All species used nearshore habitats along the coast of Argentina, with MAG also migrating further north over the Patagonian Shelf into Uruguayan waters in late autumn (May). Both SAFS and MAG have an extensive and highly abundant distribution along the Atlantic coast of South America from Argentina to Uruguay (Schiavini et al. 2005, Stokes et al. 2014, Crespo et al. 2015, Franco-Trecu 2015), with over 80 breeding and haul-out locations (refer to Fig. 1). While there are no known BBA breeding colonies in Argentina (Phillips et al. 2016), albatrosses tracked at colonies in the South Atlantic regularly use these productive coastal waters for foraging and prospecting (Croxall & Wood 2002, Clay et al. 2019). Our study provides strong evidence that BBA, SAFS and MAG from the Falkland Islands interact with conspecifics in South America, particularly in densely populated marine and terrestrial areas. Furthermore, all species were capable of transiting between South America and the Falkland Islands within 4 - 7 days, an estimate of HPAI infectious period according to experimental studies conducted on multiple gull species (Brown et al. 2008, Ramis et al. 2014). This demonstrates a substantial risk of HPAI and other pathogens spreading from South America to colonial breeding marine predator populations at the Falkland Islands (or vice versa), confirmed by the first case detection in a BBA breeding locally in late November 2023 (Falkland Islands Government 2023). While the first case of HPAI detected at the Falkland Islands was in a vagrant Southern fulmar (*Fulmarus glacialoides*) (Bennison et al. 2023), breeding BBA returning to colonies are most likely to initiate a locally transmitted outbreak, despite not visiting colonies in South America (or spending time on land). This suggests that transmission is occurring at-sea in areas of high inter-species mixing or from scavenging dead infected animals.

This study indicates an expansive spatial network and numerous pathways for HPAI to spread from the Atlantic coast of South America to the Falkland Islands. In particular, Staten Island in southern Argentina may serve as a hotspot for disease transmission. Both BBA and SAFS from the Falkland Islands spent substantial amounts of time in close connection with Staten Island, recording resident behaviours on/near land. Numerous individuals recorded repeated foraging trips to/from Staten Island. There are several SAFS breeding/haul-out sites at Staten Island (Crespo et al. 2015). The area is also an important foraging habitat for albatrosses and a range of other seabird species (i.e. penguins and flying seabirds) (Yorio et al. 2001, Croxall & Wood 2002, Pütz et al. 2006, Finger et al. 2023). Should HPAI reach Staten Island, inter-population mixing may increase the risk of exposure to infected individuals, heightening the risk of rapid disease transmission and potential disease spillover between SAFS, BBA and other migratory seabird species (Dewar et al. 2023, Leguia et al. 2023).

Understanding the temporal components of pathogen spread is also critical for effective disease surveillance. The dataset used in this study compiled BBA, SAFS and MAG movement information across different seasons and life history stages. Our findings demonstrate the risk of HPAI and pathogen spread to the Falkland Islands via these regional-ranging species is present throughout much of the year. It is particularly important to highlight breeding BBA and female lactating SAFS both showed evidence of connectivity with South America. During these periods, BBA and SAFS individuals conducted relatively short and frequent foraging trips to southern Argentina, regularly returning to colonies to incubate eggs or provision offspring. The incursion of HPAI or other zoonotic diseases to BBA or SAFS at the Falkland Islands during these critical life history stages, when there are large aggregations of conspecifics at colonies, may increase the likelihood of transmission (Duriez et al. 2023, Lane et al. 2023). As two species occurring in globally significant populations at the Falkland Islands, mass adult mortalities during the breeding period may have unprecedented conservation implications. Furthermore, high densities of seabirds and pinnipeds at the peak of the breeding season, including many likely immunologically naïve offspring, represent a fertile ground for the virus to transmit and persist within the marine predator community (Jeong et al. 2019).

The spatial pattern and chronology of HPAI spread in the region has been contrary to initial predictions. Given the Falkland Islands’ relatively close proximity to South America, the virus’ prior incursion into more distant regions south of the polar front has confounded biological risk assessments and underscored the complexity of animal movement and population connectivity as mechanisms of disease spread (Dewar et al. 2023). Surprisingly, the Falkland Islands have seemingly been buffered from the initial spread of HPAI in the region. For example, HPAI-related mass mortality events in sea lion and elephant seal populations in South America (occurring between January - October 2023) (Plaza et al., Dee Boersma 2023, Ulloa et al. 2023) are yet to trigger outbreaks at the Falkland Islands. While there is some evidence indicating spatial connectivity between Argentina and the Falkland Islands for these species (Baylis et al. 2017, Hindell et al. 2020, Campagna et al. 2021), it is possible the seasonal timing of migratory movements across the Patagonian Shelf may not coincide with ongoing outbreaks in South America, limiting/delaying the regional spread of pathogens. Alternatively, transit times between South American and Falkland Islands coastal waters may exceed the period individuals remain infectious. However, with numerous HPAI strains now circulating among seabird and pinniped species in South America, (Leguia et al. 2023, Ulloa et al. 2023, Bennison et al. 2023), spillover into other more mobile regional-ranging marine predators – such as BBA, SAFS and MAG, poses a significant threat. Other highly mobile species may also play a significant role in HPAI spread in the region. This is particularly relevant for non-breeding, scavenging seabirds, such as southern giant petrels (*Macronectes giganteus*), brown skuas (*Stercorarius antarcticus*) and sheathbills (Chionis albus), which can feed on the carcasses of infected individuals and rapidly migrate over large distances (Phillips et al. 2007, de Souza Petersen et al. 2017, Dewar et al. 2023, Bennison et al. 2023).

## 5 Acknowledgements

We are extremely grateful to the many colleagues who assisted with the fieldwork and animal tracking efforts underpinning this manuscript. All animal handling was conducted with approval of the Falkland Islands Environmental Committee and under permits issued by the Falkland Islands Government. This work was principally funded by the UK Government through the Darwin Plus Fund (DPLUS137 and DPLUS168) and funding from John Ellerman Foundation, Shackleton Scholarship Fund, Falkland Islands Government Environmental Studies Budget, and the ACAP small grants scheme. A.G. was supported by the Royal Society through a Newton International Fellowship (NIF\R1\211869) and the UK Government through the Darwin Plus Fund (DPLUS167). Data contributed by PC, LC and JP were via projects funded by 2020-2021 Biodiversa+ and Water JPI joint call for research projects, under the BiodivRestore ERA-NET Cofund (GA N°101003777), with the EU and the funding organisation FCT - Portugal (DivRestore/0012/2020; DOI 10.54499/JPCOFUND2/0001/2021) and also by FCT – Portugal through the projects: UIDB/50017/2020, UIDP/50017/2020 and LA/P/0094/2020, granted to CESAM; and UIDB/04292/2020, UIDP/04292/2020 granted to MARE and ISPA; LA/P/0069/2020 granted to the Associate Laboratory ARNET. The Falkland Islands Government provided funding through the Environmental Studies.

## 6 Author contributions

J.R and A.M.M.B designed the research; J.R performed the research; J.R and R.A.O analyzed the data; J.R, R.A.O, A.G and A.M.M.B wrote the paper; M.T, P.C, J.P.G and L.C contributed tracking data and provided input on manuscript drafts.

## 7 Supplementary Material

### Animal handling

All details of animal handling and tagging procedures are synthesised in Baylis et al (2019b), and where applicable, further described in source literature (refer to Table S1). In relation to the South American fur seal pup tracking data presented in this study, individuals were captured by hand or in a hoop net and manually restrained. Consistent with the details provided in Baylis et al (2019b), satellite tracking devices (Wildlife Computers TDR10-F-tags) were subsequently attached using a two-part epoxy (Devcon® 5-minute epoxy).

**Table S1.** Summary of all GPS and PTT telemetry data available for colonial breeding marine predator species tracked at the Falkland Islands. The * symbol in the % colonies tracked column indicates uncertainty regarding the total number of colonies at the Falkland Islands.

| Species | # individuals and device type | # and % colonies tracked | Date range | Data source |
| --- | --- | --- | --- | --- |
| Pinnipeds |  |  |  |  |
| South American fur seal<br>(*Arctocephalus australis*) | 74 (GPS and PTT) | 4 (33%) | 2018 - 2023 | Baylis et al (2019b); Riaz et al (2023); this study |
| South American sea lion<br>(*Otaria flavescens*) | 46 (GPS and PTT) | 3 (4%) | 2013 - 2017 | Baylis unpublished data; Baylis et al (2017) |
| Southern elephant seal<br>(*Mirounga leonine*) | 10 (PTT) | 1 (25%) | 2010 - 2011 | Galimberti and Sanvito unpublished data |
| Penguins |  |  |  |  |
| Magellanic penguin<br>(*Spheniscus magellanicus*) | 45 (PTT) | 6 (7%) | 2014 – 2016 | Baylis et al (2019b); Tierney unpublished data; this study |
| Gentoo penguin<br>(*Pygoscelis papua*) | 80 (GPS and PTT) | 5 (7%) | 2008 - 2015 | Baylis et al (2019b) |
| Rockhopper penguin (*Eudyptes chrysocome*) | 143 (GPS and PTT) | 6 (17%) | 1999 - 2014 | Dee Boersma et al (2002); Masello et al (2010); Putz et al (2018); Baylis et al (2019b) |
| King penguin<br>(*Aptenodytes patagonicus*) | 8 (PTT) | 1 (50%) | 2010 | Baylis et al (2015) |
| Flying seabirds |  |  |  |  |
| Black-browed albatross<br>(*Thalassarche melanophris*) | 341 (GPS) | 5 (42%) | 2008 - 2022 | Campioni et al (2017); Baylis et al (2019b); this study |
| Sooty shearwater<br>(*Ardenna grisea*) | 20 (GPS) | 1 (*) | 2017 | Baylis et al (2019b); Bonnet-Lebrun et al (2020) |

**Table S2.** List and location of all SAFS and MAG colonies in Argentina and Uruguay, collated from source literature (refer to *Methods* section). Colony ID numbers correspond with Fig. 1.

| Country | Location | Latitude | Longitude | Type | Colony ID |
| --- | --- | --- | --- | --- | --- |
| **SAFS** |  |  |  |  |  |
| Uruguay | Cabo Polonio | -34.35 | -53.73 | Breeding | 1 |
| Uruguay | Islote de Lobos | -35.02 | -54.87 | Breeding | 2 |
| Argentina | Mar del Plata | -38.10 | -57.55 | Haul-out | 3 |
| Argentina | Islote Lobos | -41.40 | -65.05 | Haul-out | 4 |
| Argentina | Isla Escondida | -43.72 | -65.28 | Breeding | 5 |
| Argentina | Cabo Dos Bahías | -44.93 | -65.52 | Haul-out | 6 |
| Argentina | Isla Arce | -45.00 | -65.48 | Haul-out | 7 |
| Argentina | Isla Rasa | -45.11 | -65.39 | Breeding | 8 |
| Argentina | Cabo Blanco | -47.22 | -65.73 | Haul-out | 9 |
| Argentina | Islote del Cabo | -48.25 | -66.22 | Haul-out | 10 |
| Argentina | Cabo Furneaux | -54.72 | -63.89 | Haul-out | 11 |
| Argentina | Caleta Ojeda | -54.72 | -63.81 | Haul-out | 12 |
| Argentina | Punta Leguizamo | -54.73 | -63.80 | Breeding | 13 |
| Argentina | Pta. Dorgambide and Pta. Shank | -54.73 | -63.94 | Haul-out | 14 |
| Argentina | Punta Jira | -54.76 | -63.81 | Breeding | 15 |
| Argentina | Punta Achával | -54.84 | -64.18 | Breeding | 16 |
| Argentina | Is. Barrionuevo (Is. Dampier) | -54.85 | -64.16 | Breeding | 17 |
| Argentina | Islote Les Eclaireurs Oeste | -54.87 | -68.10 | Haul-out | 18 |
| Argentina | Islote Veleros | -54.92 | -65.32 | Haul-out | 19 |
| Argentina | Islets south of Cabo Hall | -54.98 | -65.69 | Haul-out | 20 |
| Argentina | Islote Blanco | -55.06 | -66.55 | Haul-out | 21 |
| **MAG** |  |  |  |  |  |
| Argentina | Bahía San Antonio | -40.78 | -64.78 | Breeding | 22 |
| Argentina | Islote La Pastosa | -41.42 | -65.03 | Breeding | 23 |
| Argentina | Islote Redondo | -41.43 | -65.02 | Breeding | 24 |
| Argentina | Islote Pájaros | -41.45 | -65.03 | Breeding | 25 |
| Argentina | Punta Pozos | -41.57 | -65.02 | Breeding | 26 |
| Argentina | Ea. San Lorenzo | -42.08 | -63.85 | Breeding | 27 |
| Argentina | Asentamiento Oeste | -42.10 | -63.93 | Breeding | 28 |
| Argentina | Caleta Externa | -42.27 | -63.63 | Breeding | 29 |
| Argentina | Isla Segunda | -42.33 | -63.65 | Breeding | 30 |
| Argentina | Isla Primera | -42.35 | -63.62 | Breeding | 31 |
| Argentina | Islote Notable | -42.42 | -64.52 | Breeding | 32 |
| Argentina | Caleta Interna | -42.45 | -63.65 | Breeding | 33 |
| Argentina | El Pedral | -42.95 | -64.38 | Breeding | 34 |
| Argentina | Punta Clara | -43.97 | -65.25 | Breeding | 35 |
| Argentina | Punta Tombo | -44.03 | -65.18 | Breeding | 36 |
| Argentina | Punta Lobería | -44.58 | -65.37 | Breeding | 37 |
| Argentina | Isla Cumbre | -44.58 | -65.37 | Breeding | 38 |
| Argentina | Isla Blanca Mayor | -44.77 | -65.63 | Breeding | 39 |
| Argentina | Isla Moreno | -44.90 | -65.53 | Breeding | 40 |
| Argentina | Cabo Dos Bahías | -44.95 | -65.53 | Breeding | 41 |
| Argentina | Isla Arce | -45.00 | -65.48 | Breeding | 42 |
| Argentina | Península Lanaud | -45.05 | -65.58 | Breeding | 43 |
| Argentina | Isla Leones | -45.05 | -65.60 | Breeding | 44 |
| Argentina | Isla Sudoeste | -45.05 | -65.60 | Breeding | 45 |
| Argentina | Isla Buque | -45.05 | -65.62 | Breeding | 46 |
| Argentina | Isla Gaviota | -45.10 | -65.97 | Breeding | 47 |
| Argentina | Isla Tova | -45.10 | -66.00 | Breeding | 48 |
| Argentina | Isla Este | -45.12 | -65.93 | Breeding | 49 |
| Argentina | Isla Tovita | -45.12 | -65.95 | Breeding | 50 |
| Argentina | Isla Noroeste | -45.17 | -66.52 | Breeding | 51 |
| Argentina | Isla Fondo 1 | -45.17 | -66.52 | Breeding | 52 |
| Argentina | Isla Fondo 2 | -45.17 | -66.52 | Breeding | 53 |
| Argentina | Isla Viana Mayor | -45.18 | -66.40 | Breeding | 54 |
| Argentina | Isla Vernaci Este | -45.18 | -66.48 | Breeding | 55 |
| Argentina | Isla Vernaci Norte 1 | -45.18 | -66.50 | Breeding | 56 |
| Argentina | Isla Vernaci Norte 2 | -45.18 | -66.50 | Breeding | 57 |
| Argentina | Isla Sudoeste | -45.18 | -66.52 | Breeding | 58 |
| Argentina | Punta Pájaros | -46.95 | -66.85 | Breeding | 59 |
| Argentina | Isla Quiroga | -47.75 | -65.93 | Breeding | 60 |
| Argentina | Isla Larga | -47.75 | -65.93 | Breeding | 61 |
| Argentina | Isla Quinta | -47.75 | -65.93 | Breeding | 62 |
| Argentina | Isla Elena | -47.75 | -65.93 | Breeding | 63 |
| Argentina | Isla Pájaros | -47.75 | -65.97 | Breeding | 64 |
| Argentina | Islote del Cañadón del Puerto | -47.75 | -66.00 | Breeding | 65 |
| Argentina | Isla Chaffers | -47.77 | -65.87 | Breeding | 66 |
| Argentina | Islote Burlotti | -47.77 | -65.95 | Breeding | 67 |
| Argentina | Isla del Rey | -47.77 | -66.05 | Breeding | 68 |
| Argentina | Isla Pingüino | -47.90 | -65.72 | Breeding | 69 |
| Argentina | Isla Chata | -47.93 | -65.73 | Breeding | 70 |
| Argentina | Isla Schwarz | -48.07 | -65.90 | Breeding | 71 |
| Argentina | Islote Burgos | -48.08 | -65.90 | Breeding | 72 |
| Argentina | Isla Liebres | -48.10 | -65.90 | Breeding | 73 |
| Argentina | Punta Medanosa | -48.10 | -65.92 | Breeding | 74 |
| Argentina | Punta Sur | -48.12 | -65.93 | Breeding | 75 |
| Argentina | Estancia 8 de Julio | -48.12 | -66.13 | Breeding | 76 |
| Argentina | Islote del Bajío | -48.35 | -66.35 | Breeding | 77 |
| Argentina | Cabo Guardián | -48.35 | -66.35 | Breeding | 78 |
| Argentina | Isla Rasa Chica | -48.37 | -66.33 | Breeding | 79 |
| Argentina | Islote sin Nombre | -48.37 | -66.35 | Breeding | 80 |
| Argentina | Banco Cormorán | -49.27 | -67.67 | Breeding | 81 |
| Argentina | Banco Justicia | -49.28 | -67.68 | Breeding | 82 |
| Argentina | Isla Leones | -50.07 | -68.43 | Breeding | 83 |
| Argentina | Punta Entrada | -50.13 | -68.37 | Breeding | 84 |
| Argentina | Monte León | -50.37 | -68.88 | Breeding | 85 |
| Argentina | Isla Deseada | -51.57 | -69.03 | Breeding | 86 |
| Argentina | Cabo Virgenes | -52.35 | -68.38 | Breeding | 87 |
| Argentina | Cabo San Juan | -54.72 | -63.81 | Breeding | 88 |
| Argentina | Martillo | -54.91 | -67.38 | Breeding | 89 |

**Fig. S1.**
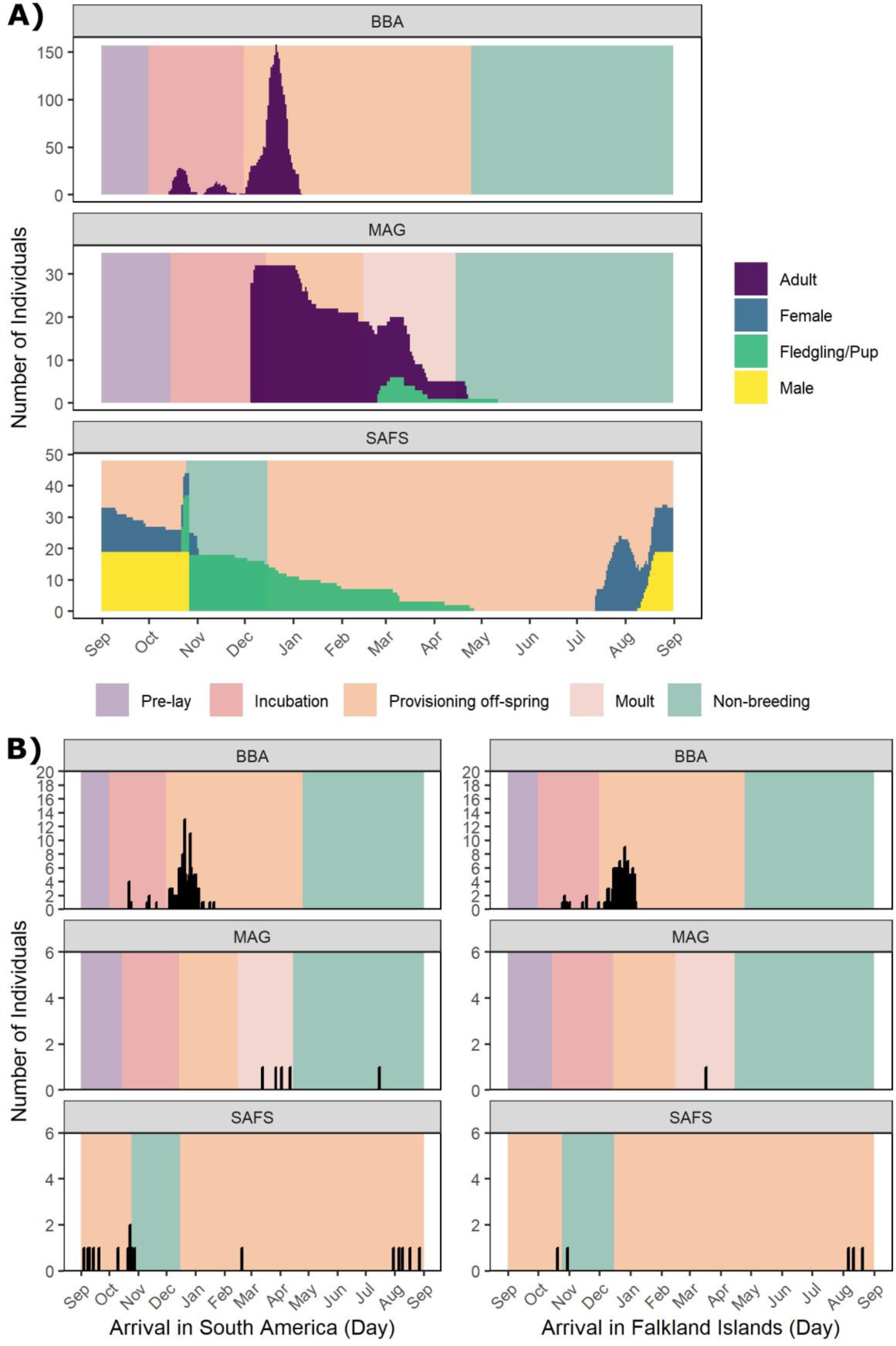
Seasonal life-history of BBA, SAFS, and MAG tracked in this study. Panels display A) all individuals tracked coloured by demographic; and B) sample size of arriving individuals between the Falkland Islands and South America, and from South America to the Falkland Islands.

**Fig. S2.**
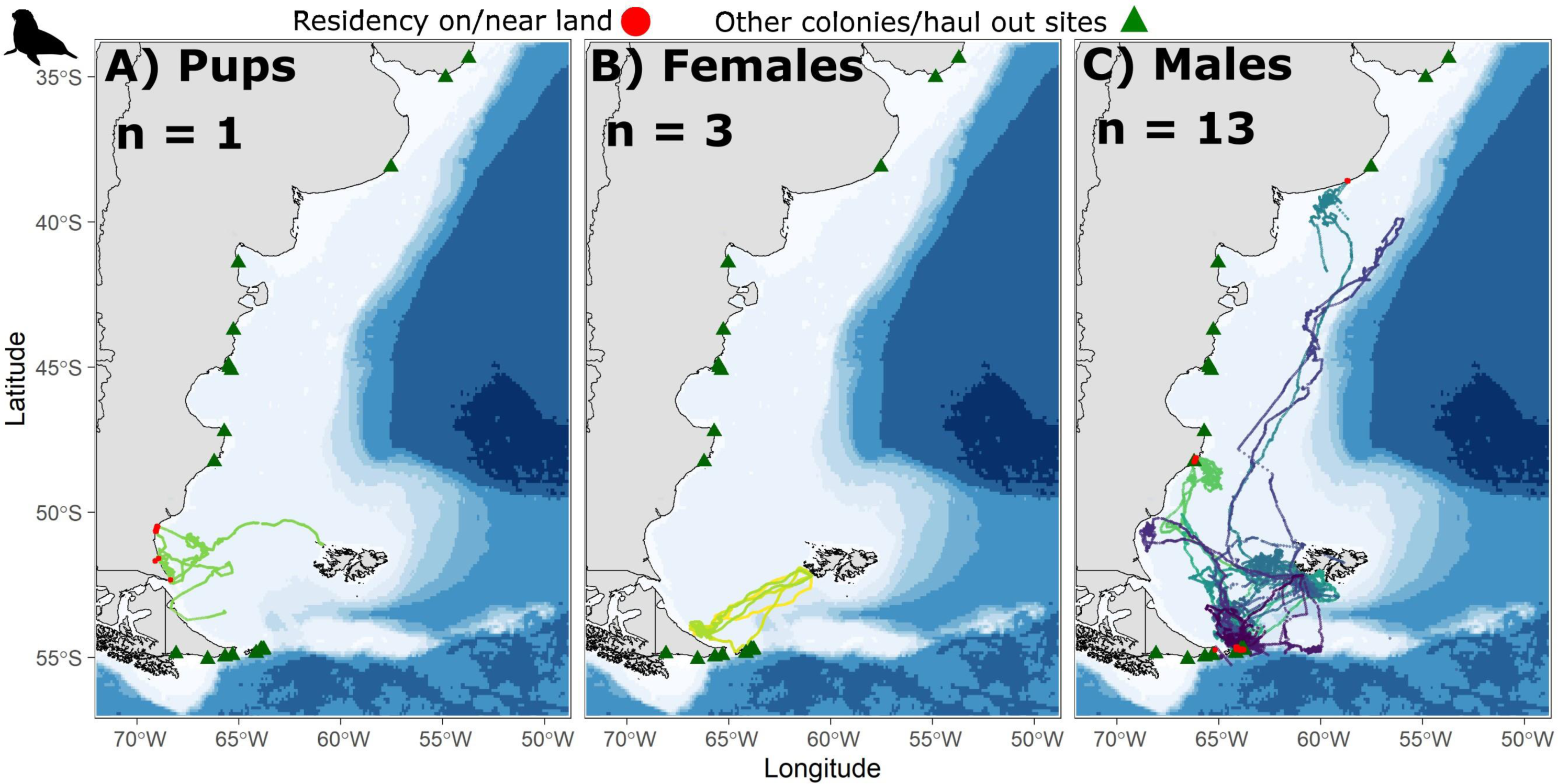
Movement data (SSM locations) of the 17 SAFS (coloured by ID and grouped by demographic) which recorded movements in coastal Argentine waters (see *Methods* for details). Map features and boundaries displayed as for Fig. 1.

## Notes

### Competing Interest Statement

The authors have declared no competing interest.

